# Timing of transcription controls the selective translation of newly synthesized mRNAs during acute environmental stress

**DOI:** 10.1101/2024.04.14.589419

**Authors:** Mostafa Zedan, Alexandra P. Schürch, Stephanie Heinrich, Pablo Aurelio Gómez García, Sarah Khawaja, Léona Dörries, Karsten Weis

## Abstract

When cells encounter environmental stress, they rapidly mount an adaptive response by switching from pro-growth to stress-responsive gene expression programs. It is poorly understood how cells selectively silence pre-existing, pro-growth transcripts, yet efficiently translate transcriptionally-induced stress mRNA, and whether these transcriptional and post-transcriptional responses are coordinated. Here, we show that following acute glucose withdrawal in S. cerevisiae, pre-existing mRNAs are not first degraded to halt protein synthesis, nor are they sequestered away in P-bodies. Rather, their translation is rapidly repressed through a sequence-independent mechanism that differentiates between mRNAs produced before and after stress followed by their decay. Transcriptional induction of endogenous transcripts and reporter mRNAs during stress is sufficient to escape translational repression, while induction prior to stress leads to repression. Our results reveal a timing-controlled coordination of the transcriptional and translational responses in the nucleus and cytoplasm ensuring a rapid and widescale reprogramming of gene expression following environmental stress.

## Introduction

The acute exposure of eukaryotic cells to environmental variations triggers rapid changes in gene expression at the transcriptional and post-transcriptional level, the so-called environmental or integrated stress response^1,2^. One hallmark of the eukaryotic stress response is the rapid inhibition of global mRNA translation to minimal levels^1^. Despite this global inhibition of translation, cells need to selectively turn on expression of genes required for adaptation to stress^2^ and ensure that these newly induced mRNAs bypass mechanisms that downregulate bulk translation^1^. How cells selectively remodel their gene expression program under stress and coordinate changes in transcription and translation remains poorly understood.

In the budding yeast *Saccharomyces cerevisiae*, acute glucose withdrawal leads to a sudden drop in ATP levels^3^ and a rapid, reversible growth arrest^4^. In this dormant state, known as quiescence, cells need to adapt their metabolism, conserve their limited energy resources and prevent the accumulation of damage until conditions improve^5,6^. To switch from a pro-growth to a stress-adaptation state, multiple layers of regulation synchronously reprogram the cell’s gene expression profile. This includes the repression of the cAMP-dependent protein kinase A and the TORC1 kinase pathways, and at the same time the induction of the AMP kinase Snf1^7^. Inhibition of the protein kinase A (PKA) and the TORC1 pathways leads to transcription inhibition of many “housekeeping” genes^7^, such as ribosomal protein and ribosome biogenesis genes, which together represent most of the cell’s transcriptome under normal growth conditions^8^. PKA inhibition also leads to the activation of the transcription factors Msn2 and Msn4, which are master regulators of the general stress response, by promoting their translocation from the cytoplasm to the nucleus^7^. Subsequently, Msn2 and Msn4 bind stress-response elements in promoters and induce the expression of hundreds of stress-responsive genes. This intricate network of transcriptional activators and repressors ensures a selective transcriptional response to acute glucose withdrawal.

Transcription regulation is, however, insufficient to rapidly switch the gene expression program upon glucose withdrawal, since at least at early time points of glucose withdrawal, a major fraction of the cellular mRNA pool is still composed of pre-existing mRNAs that were produced under glucose-rich conditions. The pre-existing mRNAs encode mostly proteins required for rapid growth and division during nutrient-rich conditions. These housekeeping mRNAs must be turned off either by mRNA degradation or translational silencing. Earlier reports have suggested that the decay rate of certain transcripts is altered following glucose withdrawal, based on analysis of mRNA levels following global transcription inhibition^9,10^. However, global transcription inhibition may result in pleiotropic effects on RNA-homeostasis^11^ or trigger feedback loops^12^, and shows weak correlation to more recent mRNA decay measurement using metabolic labeling^13–15^. A recent analysis of candidate mRNAs using metabolic labeling has shown that glucose withdrawal leads to transcript-specific effects on mRNA decay rates^16^, although the effect of glucose withdrawal on transcriptome-wide mRNA decay rates remains unclear.

In addition to changes in mRNA transcription and decay, global inhibition of translation is one of the hallmarks of the cellular response to acute glucose withdrawal^3^. This inhibition primarily targets the initiation phase of translation, leading to ribosome run-off and polysome collapse immediately after glucose withdrawal^3^. The mechanism of this translation initiation inhibition remains largely elusive, but it is independent of eIF2-alpha phosphorylation or eIF4E binding proteins and requires the loss of mRNA binding by translation initiation factors such as eIF4A, eIF4B and Ded1^17–19^. Upon acute glucose withdrawal, yeast mRNAs are also targeted to RNA-protein condensates known as P-bodies^20,21^, which are enriched for mRNA degradation factors such as Xrn1, Dcp1/2 and Dhh1. The high concentration of degradation factors initially led to the suggestion that P-bodies function in mRNA decay^20^, but deletion mutants of these factors show partial loss of translation inhibition upon glucose withdrawal, suggesting that P-bodies play a role in mRNA translation repression instead^22,23^. Furthermore, P-body purification revealed an enrichment of translationally repressed mRNAs and an absence of translation factors additionally supporting a function of P-bodies in sequestering non-translated mRNPs from the translation machinery^24^.

While bulk translation inhibition provides a means to turn off the expression of the pre-existing mRNAs, stress-induced transcripts can bypass this inhibition through mechanisms that remain largely unclear. Earlier reports have observed a tight connection between promotor sequences and translation control suggesting a crosstalk between transcription and translation regulation^25–27^. Sequence-specific mRNA elements, e.g., specific sequences in the 5′ UTR^28–30^, 3′ UTR^31^, or promoter region^25–27^, and epigenetic modifications^32,33^ have been implicated in the translation regulation of certain transcripts either by directly regulating their association with translation initiation factors^28–30^, and specific RNA binding proteins^26,31^ or through yet undefined mechanisms. It has been proposed that cells can use these mechanisms to promote the translation of certain transcripts in a targeted, sequence-specific manner. However, how hundreds of widely diverse stress-induced transcripts^2^, some of which are specific to the type of stress and are induced by very different promotors^2^ , bypass translation inhibition during stress at the global level remains poorly understood.

Here, we set out to investigate how yeast cells rapidly and selectively remodel their gene expression program following acute glucose withdrawal. We analyzed transcriptome-wide mRNA levels, mRNA decay rates and translation before and at different time points after glucose withdrawal. Our analyses reveal that cells rapidly repress the translation of mRNA produced before glucose withdrawal prior to degrading them. Efficient translation repression depends on mRNA decay factors/P-body proteins but is independent of the recruitment of mRNAs to microscopically visible P-bodies. Analysis of mRNA translation under conditions where the timing of mRNA production can be controlled - using inducible promoters or mRNA pulse labeling - demonstrates that the time of mRNA production relative to glucose withdrawal is a key determinant of mRNA translatability. The selective translation of only newly synthesized mRNA permits a facile coordination between RNA and protein production during rapid changes in gene expression.

## Results

### Accelerated decay of pro-growth mRNA following glucose withdrawal

We set out to investigate how yeast cells rapidly switch their gene expression program following acute glucose withdrawal. We first asked whether cells rapidly degrade pre-existing mRNA that was produced under glucose-rich conditions to inhibit the expression of housekeeping genes. We used SLAM-seq^34^ for pulse-chase analysis of mRNA to measure mRNA decay rates **(Fig. 1A)**. mRNA was pulse labeled using 4-thiouracil (4-TU) in glucose-rich media containing low levels of uracil to allow 4-TU incorporation below cytotoxic levels^13^, and chased with excess uracil in the presence or absence of glucose. Misincorporation of cytosine instead of thymine at alkylated 4-TU during reverse transcription enables 4-TU incorporation detection by deep sequencing at single nucleotide resolution^35^. To measure mRNA decay rates, we quantified the fraction of labeled transcripts per time point to calculate residual mRNA over time and fitted it to single phase exponential decay to calculate mRNA half-lives. Under glucose-rich conditions, 5246 transcripts could be fitted to single phase exponential decay with R-squared > 0.8 **(Fig. 1B)**, and 3283 transcripts were fitted to single phase exponential decay with R-squared > 0.8 under glucose withdrawal conditions **(Fig. 1C)**. We performed GO analysis of transcripts that were stabilized or destabilized following glucose withdrawal. While transcripts encoding mitochondrial and endosomal proteins were enriched among stabilized transcripts, transcripts encoding ribosomal proteins (RP mRNA) and ribosome biogenesis factors were highly enriched among destabilized transcripts following glucose withdrawal **(Fig. 1D)**. Analysis of residual RP mRNA over time after chase confirmed that they are long-lived under glucose-rich conditions, consistent with earlier reports^13^ but that their decay was significantly accelerated following glucose withdrawal **(Fig. 1E).** Single phase exponential decay fit per transcript showed that glucose withdrawal leads to a shift of RP mRNA half-lives from a median of 27 minutes in glucose-rich conditions to a median of 14 minutes under glucose withdrawal conditions **(Fig. 1F)**. Similar to RP mRNA, all transcripts that showed a more than 2-fold downregulation of their total mRNA levels (labeled and unlabeled reads combined) at the 30 minutes time point were significantly destabilized following glucose withdrawal when their decay rates under glucose-rich and glucose-deplete conditions were compared (total of 479 mRNAs with R-Squared > 0.8; **Fig. 1G**). These results suggest that cells employ accelerated mRNA degradation to downregulate pre-existing mRNA and adapt their gene expression program.

**Figure 1:**
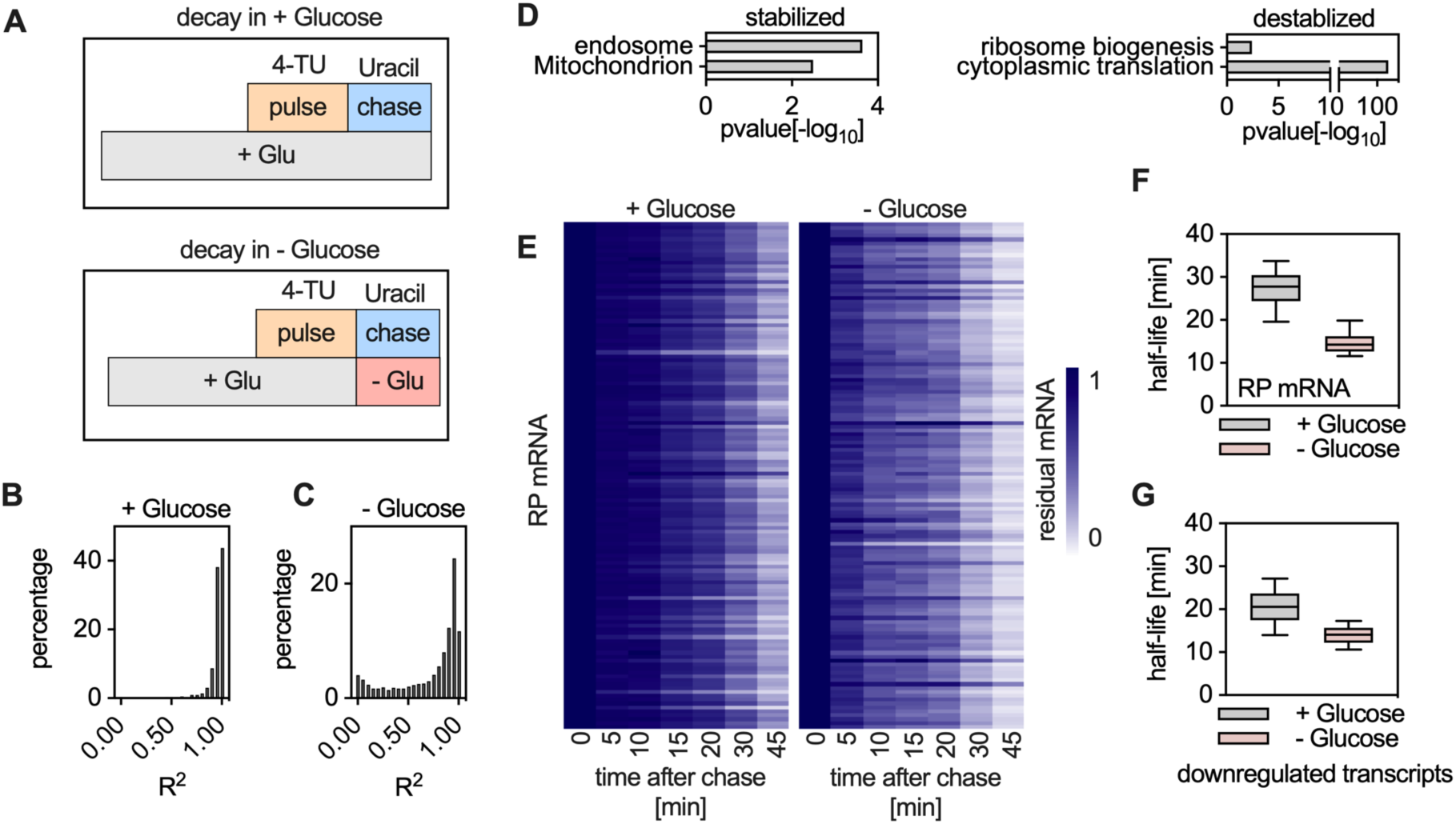
Accelerated mRNA decay following glucose withdrawal. **A.** Experimental layout of SLAM-seq analysis: Yeast cells were grown to log phase in synthetic media containing low uracil, followed by addition of 4-thiouracil to pulse label newly produced mRNA for 2h. Cells were filtered and washed to remove 4-TU and resuspended in a growth medium containing excess uracil in the presence or absence of glucose to stop RNA labeling and start the chase. Cells were collected at the start of the chase (time point zero) and after 5, 10, 15, 20, 30 and 45 min after the chase. The conversion rate was used to estimate the fraction of labeled transcripts during the chase under glucose-rich conditions. To calculate the fraction of labeled transcripts under glucose withdrawal conditions, reads with T>C conversions were normalized to in vitro transcribed RNA that was spiked-in at a constant ratio to total RNA per time point. Residual mRNA was calculated as a ratio of the fraction of labeled transcripts per time point to time point zero at the beginning of the chase. The experiment was done in two replicates and the average of the two replicates was used for the analysis. **B and C.** Histogram showing the distribution of the calculated R-Squared values for the single-phase exponential decay fitting per transcript during the chase under glucose-rich conditions (**B**), or under glucose-withdrawal conditions (**C**). **D.** GO analysis of stabilized or destabilized transcripts following glucose withdrawal, based on the calculated half-life per transcript. Only GO terms with more than 10 genes are shown. **E.** Heat map showing residual mRNA per transcript encoding ribosomal proteins during the chase under glucose-rich or glucose withdrawal conditions. **F.** Box plot showing the distribution of half-lives of transcripts encoding ribosomal proteins, with calculated R-Squared > 0.8, under glucose-rich or glucose withdrawal conditions. **G.** Box plot showing the distribution of half-lives under glucose-rich or glucose withdrawal conditions for 479 transcripts whose levels are > 2-fold downregulated after 30 minutes of glucose withdrawal. Only transcripts with calculated R-Squared > 0.8 were included in the analysis.

### Translation repression precedes mRNA decay following glucose withdrawal

Although cells accelerate decay of pre-stress housekeeping mRNA following glucose withdrawal, the changes in mRNA levels are slow. For example, based on our half-life measurements, the pool of pre-existing, ‘old’ RP mRNA is only reduced by 50% in ∼ 15 minutes. This is in contrast to changes in mRNA translation, which are extremely rapid and can occur as early as 30 seconds following glucose starvation^19^. The fast drop in mRNA translation thus cannot be explained only by mRNA decay and suggests additional regulation at the level of translation. To analyze transcriptome-wide changes in translation independent of mRNA copy number, we switched yeast cells from glucose-rich media to media that lacks glucose using rapid filtration, and harvested cells after a total of 5, 15 and 30 minutes of glucose withdrawal for a time-resolved analysis of the cellular response to glucose withdrawal. We then split the samples in two and performed RNA-seq and ribosome profiling^36^ in parallel using the same library preparation procedure. This enabled a direct comparison of total and translating levels of mRNA and allowed us to calculate the translation efficiency per transcript. Differential gene expression analysis showed that glucose withdrawal leads to the upregulation of hundreds of transcripts at both mRNA copy number and ribosome footprint levels after 5 minutes, and that this becomes more pronounced after 15 and 30 minutes **(Fig. 2A)**. GO analysis **(Fig. 2B)** of mRNA with >2-fold upregulation at the translation level following 5 minutes of glucose withdrawal showed that the upregulated genes were enriched for genes encoding stress proteins, including those involved in protein folding, such as chaperones, and proteins required for cell wall organization.

**Figure 2:**
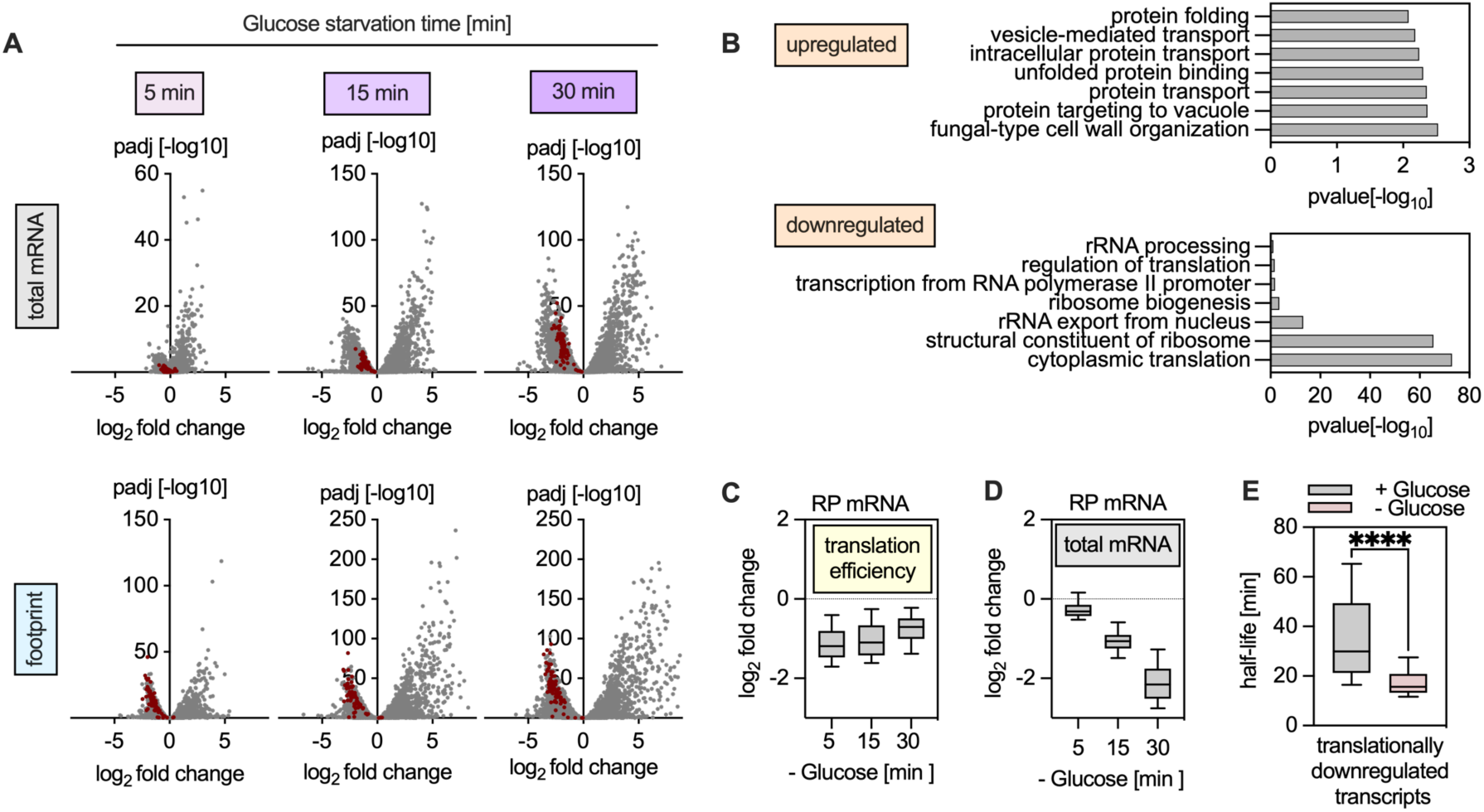
Translation repression precedes mRNA decay following glucose withdrawal. Glucose was removed from the medium by shifting yeast cells growing in glucose-rich media to glucose-free media by rapid filtration for a total of 5, 15, or 30 minutes. The level of translation was quantified by sequencing of the ribosome footprints on the mRNA using ribosome profiling. Total RNA levels were quantified by sequencing poly(A) mRNA in parallel. The experiment was done in two replicates and the average of the two replicates was used for the analysis. **A.** Volcano plots showing differentially expressed transcripts using Deseq2 at the total mRNA levels (top) as well as the ribosome footprint levels (bottom) for all detected mRNAs, after 5, 15 or 30 minutes of glucose withdrawal. mRNAs encoding ribosomal proteins (RP mRNAs) are highlighted in red. **B.** GO analysis using DAVID of mRNAs with > 2-fold up- or down-regulation at the translation level following 5 minutes of glucose withdrawal, as compared to glucose-rich conditions. Only the top enriched GO terms including more than 10 genes are shown. **C.** Box plot showing the translation efficiency of RP mRNAs following glucose withdrawal. Translation efficiency was calculated by normalizing ribosome footprint levels to total mRNA levels. Whiskers represent the 10-90% interval. **D.** Box plot showing total RP mRNA levels changes in log_2_ following glucose withdrawal after 5, 15 and 30 minutes of glucose withdrawal. Whiskers represent the 10-90% interval. **E.** Comparison of mRNA half-lives under glucose withdrawal versus glucose-rich conditions, for transcripts that are 2-fold downregulated at the translation level following 5 minutes of glucose withdrawal.

Consistent with our measured mRNA decay rate kinetics, 5 minutes of glucose withdrawal led to barely detectable downregulation of any transcript at the mRNA copy number yet resulted in downregulation of many transcripts at the mRNA translation level **(Fig. 2A).** After 15 and 30 minutes of glucose withdrawal, many transcripts were downregulated at both the mRNA copy number and translation levels based on footprints **(Fig. 2A).** GO analysis showed that genes which were downregulated more than 2-fold at the translation level following 5 minutes of glucose withdrawal were highly enriched in housekeeping genes, such as those involved in mRNA translation and ribosome biogenesis **(Fig. 2B)**. mRNAs encoding ribosomal proteins were the group most significantly downregulated at the level of translation **(Fig. 2A, 2B)**. Analysis of the translation efficiency of RP mRNA by direct comparison of their ribosome footprint levels and total mRNA levels confirmed that the translation efficiency of RP mRNA was already maximally downregulated following 5 minutes of glucose withdrawal and persisted over prolonged glucose withdrawal to 15 or 30 minutes **(Fig. 2C),** demonstrating a rapid and sustained translation repression mechanism for these mRNAs. Although RP mRNA levels were nearly unaltered after 5 minutes of glucose withdrawal, RP mRNA levels decreased after prolonging glucose withdrawal to 15 and 30 minutes **(Fig. 2D).** While this shows that mRNA decay contributes to the downregulation of the expression of these genes, it trails the effects on translation repression. Comparison of the decay rates in glucose-rich and glucose-depleted conditions of all 121 transcripts that were translationally repressed by more than 2-fold demonstrates that this entire group of transcripts is destabilized following glucose withdrawal **(Fig. 2E)**. This further confirms that translation repression precedes mRNA degradation.

### Translation repressed mRNAs are not enriched in P-bodies

Earlier reports have shown that mRNA decay factors such as Dhh1 and Pat1 can act as general repressors of translation during glucose starvation^37,38^. These decay factors are enriched together with mRNA in P-bodies, biomolecular condensates that become microscopically detectable upon acute glucose withdrawal in yeast. Since cells lacking these mRNA decay factors are also compromised for P-body formation, we asked whether the translation repression is mediated by mRNA sequestration into P-bodies. We focused on the group of mRNAs encoding ribosomal proteins since they are the most enriched among translationally repressed transcripts. Consistent with this hypothesis, ribosome profiling showed that deletion of *DHH1* and to a lesser extent also deletion of *PAT1* impaired the repression of RP mRNA translation upon glucose withdrawal **(Fig. 3A)**. Since deletion mutants of major decay factors such as Pat1 and Dhh1 can have pleiotropic effects beyond P-body formation, we also performed gene expression analysis in cells expressing only a truncated version of Dhh1 lacking the N- and C-terminal tails (Dhh1^tailless^), which reduces P-body formation without affecting Dhh1 core domains^39^. Interestingly, the observed translation repression upon glucose starvation was also significantly impaired in Dhh1^tailless^ mutant cells and was comparable to cells entirely lacking Dhh1 **(Fig. 3A)**.

**Figure 3:**
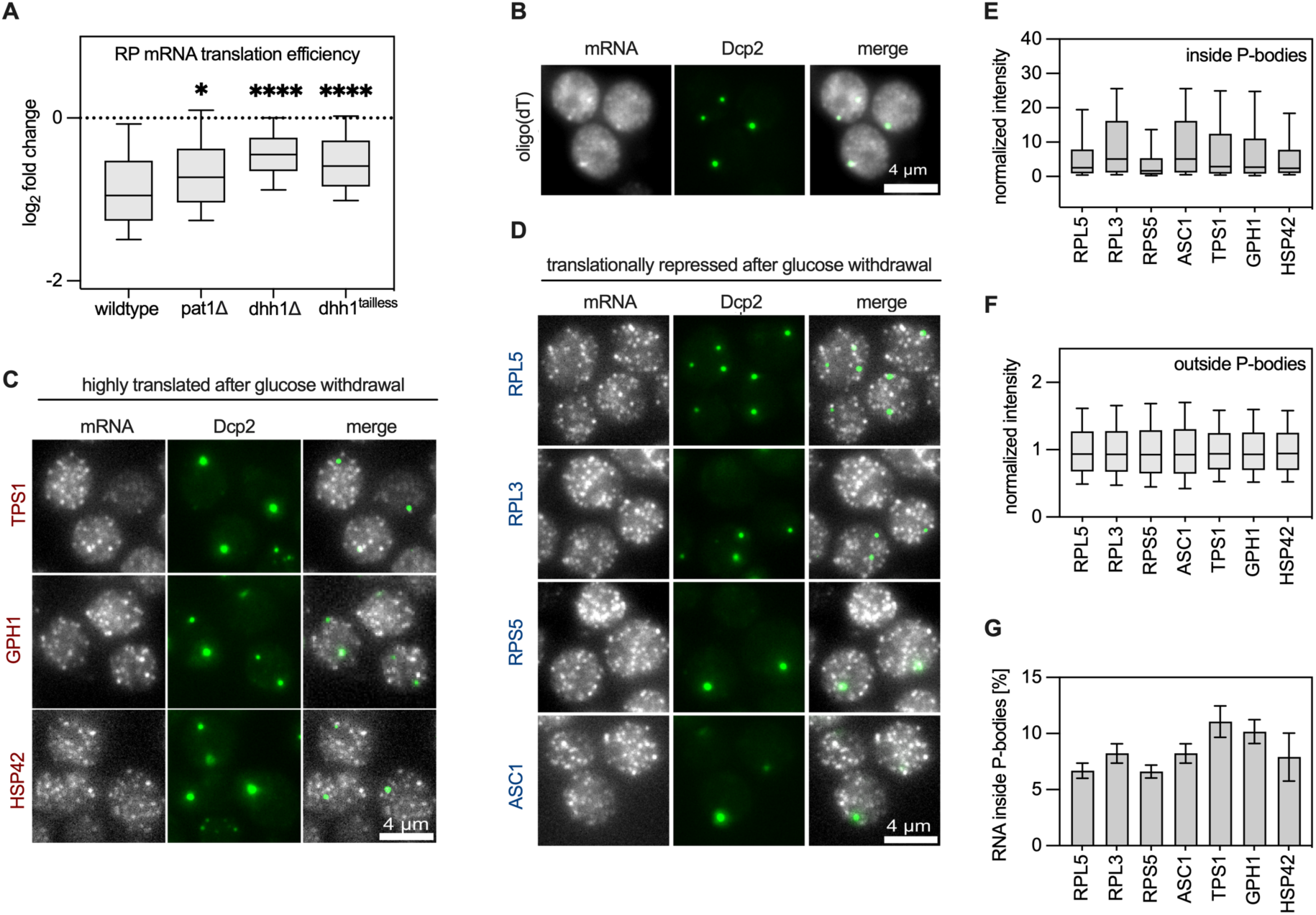
Translationally repressed mRNAs predominantly localize outside P-bodies. **A.** Box plot showing RP mRNA translation efficiency changes in log_2_ following glucose withdrawal after 5 minutes of glucose withdrawal in wild type yeast cells, or cells lacking Pat1, or Dhh1 or cells expressing only a truncated Dhh1 variant that lacks both its N- and C-terminal tails (Dhh1^tailless^). Translation efficiency was calculated by normalizing ribosome footprint levels to total mRNA levels. The experiment was done in replicates and the average of two replicates was used for the analysis. **B.** smiFISH analysis of poly(A) mRNA using oligo(dT) primary probes. Glucose was removed from the medium by shifting yeast cells expressing GFP tagged Dcp2 at the endogenous locus (as a marker of P-bodies) from glucose-rich medium to glucose-free medium by rapid filtration and incubation for 30 minutes. The oligo(dT) primary probe containing the flap sequence was first hybridized to a double labeled oligo with TAMRA dye (secondary probe) complementary to the flap-sequence of the primary probes, before it was hybridized to the target mRNA (poly(A) mRNA). **C and D.** Similar to B, except that mRNA-specific primary probes were used. 27-48 primary probes per target mRNA were used to localize translationally upregulated **(C)** or translationally downregulated **(D)** mRNAs following glucose withdrawal for 30 min. Translationally downregulated mRNAs following glucose withdrawal are colored in blue, and translationally upregulated mRNAs following glucose withdrawal are colored in red. **E.** Analysis of the fluorescence intensity per mRNA focus, detected inside P-bodies, normalized to the median of the fluorescence intensities of all detected mRNA foci. **F.** Similar to E, but only mRNAs localizing outside of P-bodies were included in the analysis. **G.** Analysis of the fraction of mRNAs that localize to P-bodies by comparing the total fluorescence intensity of detected mRNA inside P-bodies as a fraction of the total fluorescence intensities of all detected mRNA.

To directly test whether the translation repression of mRNA in glucose starvation is mediated by sequestration of these mRNAs inside P-bodies, we performed mRNA localization experiments using the smiFISH protocol^40^. In smiFISH, primary probes complementary to target mRNA sequences contain an additional flap sequence at the 5′ ends that is used for labeling and detection via a fluorescently labeled secondary probe. To localize bulk mRNA following glucose withdrawal, we used an oligo(dT) primary probe to target the poly(A) tails of mRNA. Consistent with earlier reports^41^, we detected only a weak enrichment of the probe inside P-bodies following glucose withdrawal **(Fig. 3B)**, indicating that although a fraction of total mRNA is deposited in P-bodies, the majority of the poly(A) mRNA remains outside P-bodies. To test the hypothesis that translationally repressed mRNA is enriched in P-bodies while translationally active mRNA remains outside P-bodies, we selected several mRNAs that were either translationally up- or downregulated following glucose withdrawal and determined their subcellular localization via smiFISH. To this end, multiple (27 to 48) primary probes complementary to different segments of the target mRNA sequence were used. The hybridization of multiple fluorescently labeled probes per mRNA amplified the signal-to-noise ratio, enabling single molecule mRNA visualization^40^. We filtered candidate mRNAs for expression levels and length, to ensure high signal-to-noise ratio for single molecule detection, and for the absence of sequences with high similarity, to avoid mistargeting. We performed smiFISH in a strain expressing a GFP-tagged version of the P-body marker Dcp2, in the presence of glucose or 30 minutes after glucose withdrawal, since P-bodies are prominent at this time point, and we still observed strong translation repression **(Fig. 2C)**. All tested mRNAs showed numerous distinct fluorescent foci with uniform intensity, indicative of single molecule resolution in both glucose-rich or glucose withdrawal conditions **(Supplementary Fig. 1A**). Repeating the smiFISH using only the labeled secondary probes in the absence of primary probes resulted in loss of fluorescence signal, confirming the specificity of the smiFISH analysis **(Supplementary Fig. 1B**). Interestingly, following glucose withdrawal, the large majority of fluorescent foci for both translationally upregulated **(Supplementary Fig. 1A**, **Fig. 3C)** or translationally downregulated **(Supplementary Fig. 1A**, **Fig. 3D)** mRNA remained cytoplasmic and did not colocalize to P-bodies. Due to the diffraction limits of fluorescence microscopy, mRNA spots that co-localize (i.e., that are less than ∼ 200 nm apart) cannot be differentiated. Thus, sequestration of multiple mRNAs in one P-body will not lead to an increase in the number of detectable spots in a P-body but instead will lead to an increase in RNA spot fluorescence intensity. Therefore, we also systematically quantified the signal intensity of individual mRNA foci inside and outside of P-bodies, as well as the fraction of mRNA that localize to P-bodies for each of the tested candidates. Our analysis showed that mRNA inside P-bodies tend to have an overall higher and more heterogeneous fluorescence signal **(Fig. 3E)** as compared to mRNA detected outside P-bodies **(Fig. 3F)**, indicating that multiple mRNA molecules of the same targeted mRNA can indeed cluster in P-bodies, consistent with poly(A) mRNA enrichment inside P-bodies **(Fig. 3B)**. However, neither mRNAs that were translationally downregulated following glucose withdrawal (RPL5, RPL3, ASC1 and RPS5) nor those that were translationally upregulated (TPS1, GPH1 and HSP42) showed enrichment in P-bodies, and only a minor fraction of the signal (5-10%) localized to Dcp2-positive granules **(Fig. 3G).** This shows that while P-bodies contain mRNA, permanent sequestration of an mRNA to P-bodies is not required to alter translation efficiency suggesting that translational repression can largely occur outside of P-bodies.

### The timing of mRNA production determines translation efficiency

Despite the observed overall translation repression, a significant number of mRNAs are upregulated both at the level of transcription and translation following glucose withdrawal. Analysis of differentially expressed genes following glucose removal showed a positive correlation between total mRNA levels and mRNA translation **(Fig. 4A)**. This suggests that transcripts that are transcriptionally induced after glucose withdrawal can escape translation repression, in contrast to the transcriptionally repressed, pre-existing transcripts, which are targets of rapid translational repression and subsequent degradation. Analysis of the levels of stress-induced transcripts and their translation efficiency following glucose withdrawal **(Fig. 4B)** showed that while translation repression for pre-existing transcripts occurs promptly and persists after stress, the transcription of stress-induced transcripts and their translation continues during acute glucose starvation with a constant translation efficiency.

**Figure 4:**
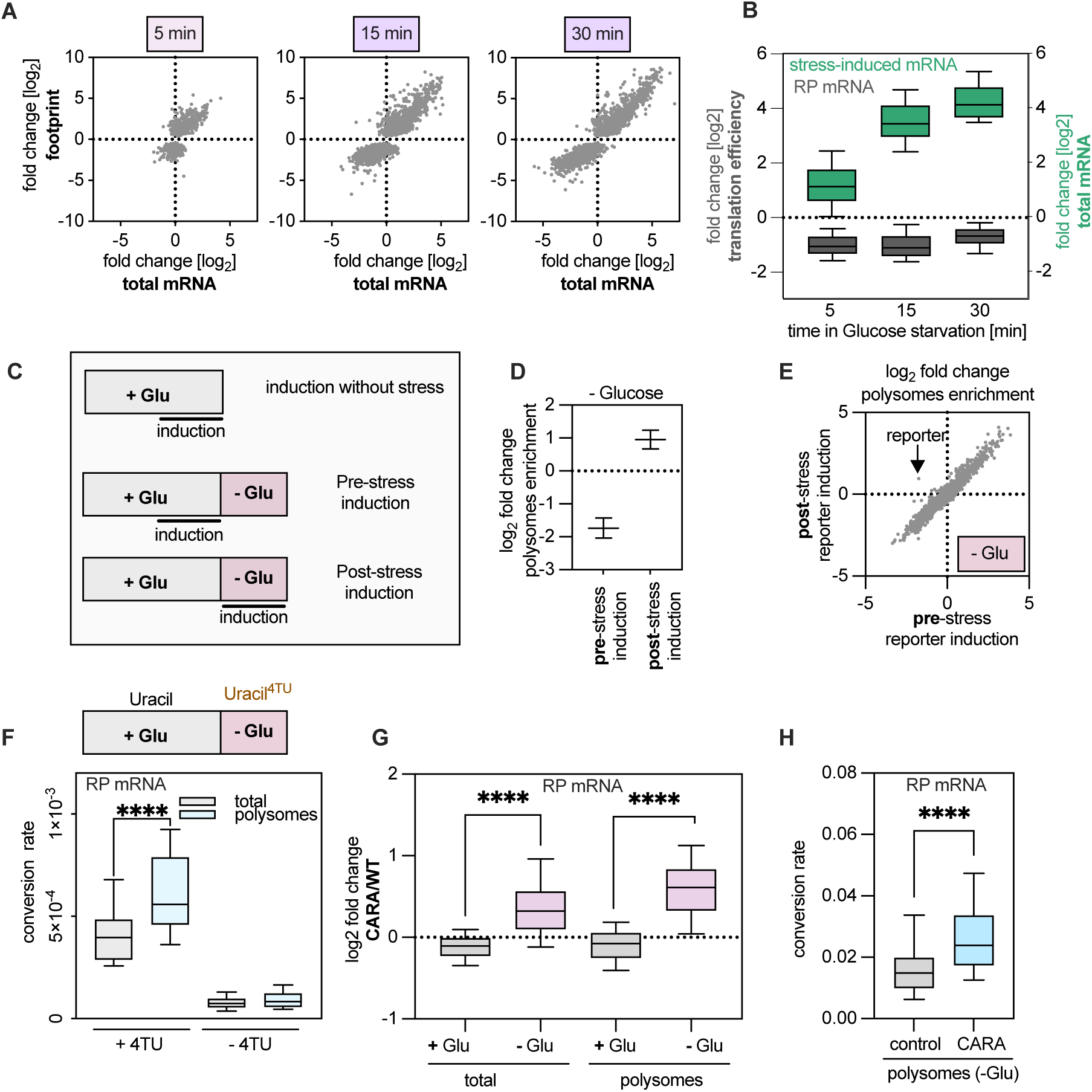
The timing of mRNA production determines translation efficiency. **A.** Correlation of the changes in mRNA translation and total mRNA levels in log_2_ for differentially translated transcripts, based on Deseq2 analysis, at 5, 15 or 30 minutes after glucose withdrawal. **B.** Box plot showing the fold change (in log_2_) of total mRNA levels or translation efficiency at 5, 15 or 30 minutes after glucose withdrawal compared to glucose-rich conditions, for transcripts that are more than 2-fold induced after 30 minutes of glucose starvation. Translation efficiency was calculated by normalizing ribosome footprint levels to total mRNA levels. **C**. Layout of the different induction schemes of an estradiol-inducible GST reporter mRNA. For the control induction *without stress*, the reporter mRNA production was induced under glucose-rich conditions (SCD) by a 30-minute estradiol addition prior to harvesting. For *pre-stress* induction, the reporter mRNA was induced under glucose-rich conditions (SCD), followed by rapid glucose withdrawal and termination of induction by filtration of yeast cells from glucose-rich medium to glucose-free medium lacking estradiol, followed by a 30-minute incubation before harvesting. For *post-stress* induction, the reporter mRNA production was induced only upon glucose withdrawal. Glucose was removed and reporter expression was induced by shifting yeast cells from glucose-rich medium to glucose-free medium containing estradiol and incubated for 30 minutes prior to harvesting. To quantify mRNA translation, the cell lysate was fractionated using a sucrose gradient followed by polysome isolation, RNA extraction and sequencing. Total RNA levels were quantified by sequencing poly(A) mRNA. **D.** Analysis of the change in polysome enrichment (in log_2_) of an estradiol-inducible reporter mRNA encoding GST following glucose withdrawal under pre- or post- mRNA induction conditions, compared to glucose-rich conditions under which the reporter was also induced. The enrichment of mRNA in polysomes was quantified by normalizing mRNA levels in the polysome fraction to total levels. **E.** Analysis of the changes in polysome enrichment (in log_2_) of all quantified mRNA following 30-minute glucose withdrawal under pre-stress or post-stress reporter induction conditions, as compared to growth in glucose- rich conditions under which the reporter was also induced. The mRNA enrichment in the polysomes was quantified by normalizing mRNA levels in the polysome fraction to its total levels. **F.** Analysis of polysome enrichment of newly made transcripts encoding ribosomal proteins after glucose withdrawal. To label newly made transcripts after glucose withdrawal, yeast cells were shifted from glucose-rich medium to glucose-free medium lacking uracil but containing 4-TU for 30 minutes before harvesting. To analyze the level of mRNA translation, cell lysates were fractionated using a sucrose gradient followed by polysome isolation. Labeled transcripts in the total lysate or in the polysomes were quantified based on the calculated T>C conversion rate. The experiment was done in two replicates. To control for background T>C conversion, the experiment was repeated without addition of 4-TU. **G**. Box plot showing the change of RP mRNA levels between the CARA and wild type strains (in log_2_), under glucose rich conditions or 30 minutes of glucose starvation, in the total lysate or in the polysomes fraction. The experiment was performed in two replicates. **H.** Comparison of the levels of post-stress RP mRNA in the polysome fraction between the control and CARA strains. To label newly made transcripts after glucose withdrawal, yeast cells were shifted from glucose-rich medium to glucose-free medium lacking uracil but containing 4-TU for 30 minutes before harvesting. To analyze mRNA translation levels, cell lysates were fractionated using a sucrose gradient followed by polysome isolation. Labeled transcripts in the polysomes were quantified based on the calculated T>C conversion rate. The experiment was performed in two replicates.

We therefore asked whether the timing of mRNA production relative to glucose withdrawal determines the mRNA translation fate. To address this question, we took advantage of an estradiol-inducible promoter system^42^ which allowed us to control the time of production of a reporter mRNA expressing the GST protein **(Fig. 4C)**. For pre-stress induction of the mRNA, we induced the transcription of the reporter in the presence of glucose before shifting cells to media lacking glucose and containing no estradiol. To mimic a stress-induced mRNA, we induced the reporter at the time of glucose withdrawal. As a control, we also induced transcription of the reporter in the sustained presence of glucose. To measure mRNA translation in each of the different induction programs, we compared mRNA levels in isolated polysomal fractions to total mRNA levels in the lysate. Consistent with a global translation repression upon glucose withdrawal, the reporter was less enriched in polysomes when it was induced prior to glucose stress **(Fig. 4D)**. Remarkably, the identical reporter mRNA when induced during stress escaped translational repression and was highly associated with polysomes after glucose starvation **(Fig. 4D)**. Importantly, the addition of estradiol did not affect the regulation of endogenous transcripts, and global changes in polysomal enrichment following glucose withdrawal were indistinguishable between conditions where the reporter was induced pre- or post-stress **(Fig. 4E)**. Similar analysis of an inducible reporter mRNA encoding citrine also showed translation regulation depending on the timing of transcription before or after glucose withdrawal **(Supplementary Fig. 2A),** without affecting the repression of RP mRNAs following depletion of glucose **(Supplementary Fig. 2B**). Thus, differential repression can be observed with identical transcripts driven by the same promoter. This reveals a sequence-independent mechanism of translation regulation that is exclusively dependent on the timing of transcription.

We next asked whether endogenous mRNA transcribed from the same gene before and during stress would also experience different translation fates. To address this question, we pulse-labeled newly made transcripts during stress by shifting cells from media containing glucose and uracil to media lacking glucose and containing 4-TU to exclusively label newly made transcripts during glucose withdrawal. We then compared the levels of 4-TU incorporation per transcript for polysome-associated mRNA as well as total mRNA extracted from the total lysate. Overall, we observed low levels of 4-TU incorporation following glucose removal, consistent with a global transcriptional reduction upon glucose withdrawal^43^. The production of RP mRNAs, which we found to be strongly translationally repressed during glucose starvation **(Fig. 2C)**, was also clearly repressed at the transcriptional level. However, their transcription was not completely abrogated, and RP mRNAs still showed low levels of 4-TU incorporation following glucose withdrawal **(Fig. 4F)**. The detected incorporation rate was significantly higher than background, where no 4-TU was added **(Fig. 4F)**, allowing us to ask whether the residual RP mRNA produced after stress would escape translation repression. Strikingly, the newly made, 4-TU-marked RP mRNA that was produced after glucose withdrawal could be found at higher levels in the polysomes compared to total mRNA. Furthermore, while the newly made RP mRNA was enriched in polysomes and thus escaped translational repression **(Fig. 4F)**, the bulk of the RP mRNA, which predominantly consists of mRNAs produced prior to stress, was still translationally repressed under these conditions **(Supplementary Fig. 2C**), consistent with our ribosome profiling analysis **(Fig. 2C).** Similar to RP mRNA, newly made transcripts for other mRNAs that are translationally repressed in bulk under acute glucose stress were enriched in polysomes, further confirming that mRNAs produced before and after stress have different translation fates **(Supplementary Fig. 2D**). These results demonstrate that for endogenous mRNA too, the timing of production is a key determinant of translation fate during acute glucose stress, independent of promoter or transcript sequence.

To investigate whether the preferred use of newly made transcripts is a general feature of the translation machinery, we also performed pulse-chase experiments under glucose-rich conditions. RNA was labeled with a short pulse of 4- TU followed by a chase with excess uracil. The levels of labeled RP mRNAs decreased steadily between 0, 10 and 20 minutes after the addition of excess uracil, confirming an efficient chase of RP-mRNA labeling **(Supplementary Fig. 2E**). To estimate translation efficiency, we analyzed the levels of labeled mRNA in the total and in the polysomes fraction. Interestingly, the labeled mRNA showed a similar translation efficiency at all time points of the experiment **(Supplementary Fig. 2F**). This suggests that newly-made transcripts are not preferentially translated under glucose-rich conditions, and the selection of new over old mRNAs is specific for translation upon stress induction.

Our results predict that a perturbation of the timing of mRNA production would affect translation control during glucose stress. We tested this by taking advantage of the CARA (Constitutive Association of Rrn3 and A43) yeast strain^44^. This strain expresses a constitutively active form of RNA polymerase I (pol- I) preventing the repression of pol-I transcription during nutrient stress. As a consequence, CARA cells also fail to properly repress transcription of mRNAs encoding ribosomal protein upon glucose withdrawal. We therefore asked whether RP-mRNA production and translation is elevated in the CARA strain in glucose starvation conditions. The levels of RP-mRNAs were compared in the total lysate as well as the polysomes fraction between wild type and CARA cells before and after glucose starvation. In the presence of glucose, RP-mRNA levels were comparable in the wild type and CARA strain. However, the CARA strain showed higher levels of RP mRNA upon glucose starvation, consistent with a failure to efficiently repress the production of RP mRNA during stress. Importantly, the CARA strain also displayed higher levels of RP mRNA in the polysomes fraction following glucose starvation, indicating that the RP mRNAs that are produced during glucose withdrawal escape translation repression **(Fig. 4G)**. To directly confirm this, we also performed RNA labeling using 4-TU during glucose starvation in the CARA strain and determined the amount of labeled mRNA in the polysomes fraction. Interestingly, higher levels of labeled, post-stress transcribed RP mRNA were found in the polysomes fraction of CARA cells compared to the wild-type strain. These results confirm that newly produced mRNA selectively escapes the global translation repression upon glucose withdrawal, further demonstrating that cells employ a timing-controlled and sequence-independent mechanism to establish selectivity of its translation program upon acute stress.

## Discussion

Organisms frequently encounter environmental challenges that require them to mount an adaptive response by changing their gene expression program. A rapid and wholescale reorganization of the cell’s transcriptional and translational activity is seen in diverse stresses such as temperature shifts, osmotic shock, hypoxic conditions and nutrient deprivation, and this environmental or integrated stress response is crucial for cell survival^2^. In *S. cerevisiae*, glucose is the preferred energy source and an important signaling molecule that regulates the cell’s gene expression program through multiple signaling pathways^7^. At the transcriptional level, upon acute glucose withdrawal, yeast cells downregulate a large number of genes that are required for rapid growth and proliferation (referred to as “*housekeeping”* genes) and induce stress-related genes as part of an adaptive response that allows them to enter a state of quiescence and change their energy metabolism^18^. Adaptation needs to occur fast, yet at early time points of glucose withdrawal, most of the transcriptome is still composed of mRNA produced under glucose-rich conditions, largely encoding housekeeping proteins. To halt energy-consuming and growth-promoting processes, expression of these housekeeping genes must thus be swiftly inhibited by post-transcriptional mechanisms. Here, we show that translation repression and mRNA decay converge to inhibit the expression of pro-growth, pre-stress transcripts following glucose withdrawal in a time-resolved fashion. Following glucose withdrawal, we observe a very rapid and selective reduction in the translation efficiency of housekeeping transcripts, independent of mRNA copy number, that reaches its maximal levels already at 5 minutes, the earliest time point that we measured, consistent with the earlier reported very rapid displacement of initiation factors upon glucose withdrawal^19^. This translation repression is then temporally succeeded by mRNA decay, which is accelerated for mRNAs that are translationally repressed, enabling cells to gradually downregulate the levels of these pre-stress transcripts. These results suggest a coordination between mRNA translation and degradation which is in line with many experiments that have demonstrated a tight connection between these two processes (e.g., ^13,45–47^). While the mechanism and effects of this coupling are still debated, our data are consistent with a tug-of-war model between the translation and the decay machineries, which postulates that actively translated transcripts are protected against decay and are therefore long-lived, whereas translationally repressed mRNAs undergo accelerated decay.

Our ribosome profiling analyses highlighted another link between the regulation of mRNA translation and decay, demonstrating that the critical decay and P-body factors Dhh1 and Pat1 are important for the efficient repression of housekeeping mRNA translation upon glucose withdrawal. These results also validate the findings that mutants of Dhh1, Pat1 and other mRNA decay factors display diminished polysome collapse upon glucose withdrawal^37,38^. Furthermore, we find that a mutant of Dhh1 that lacks its unstructured C- and N-terminal tails also shows impaired translation repression upon glucose withdrawal. Since the tails of Dhh1 play an important role in P-body assembly but don’t affect Dhh1’s core domains^39^, these results could suggest that P-body formation is important for translational repression or that cells sequester mRNA in a translationally inactive state in P-bodies. However, our RNA localization experiments speak against both large-scale and selective storage of repressed mRNA in P-bodies. First, we found less than 10% of any tested, repressed mRNA in a P-body 30 minutes after glucose withdrawal (Fig. 3). Thus, repressed mRNPs are not sequestered into P-bodies *en masse* during stress. Second, we observed maximal translation repression already after 5 minutes of glucose starvation, when we do not yet observe large microscopically detectable P-bodies (Supplementary Fig. 1C). Nevertheless, it is likely that translation repression requires multimerization of decay factors such as Dhh1 on the mRNA. This is probably at least partially mediated by the unstructured tails of Dhh1 forming repressive complexes that keep the mRNA in a translationally inactive state. Our results would be consistent with a model that repressed mRNPs would form in the cytoplasm and then condense over time into microscopically detectable P bodies. It should be noted that our methods cannot address whether rapidly forming, repressive RNA-protein complexes contain only a single mRNA or whether several, distinct mRNAs can come together to form ‘micro-condensates’ or ‘micro-P-bodies’. However, our results are inconsistent with the hypothesis that rapid mRNA deposition into one or a few large P-bodies is required for translation repression. In this model, the formation of large, microscopically detectable P-body condensates would be a consequence but not a cause of bulk translational repression.

Earlier reports have used stem-loop RNA labeling systems to visualize single molecules of mRNA in live cells^48^ which has led to the notion that P-bodies can be sites of translation repression^25,49^. However, these systems in their original implementation may have resulted in an overestimation of the RNA fraction inside P-bodies, since stem-loop insertion can lead to the accumulation of degradation intermediates^50^. Furthermore, single, stem-loop labeled mRNPs are dim and diffuse fast in living cells^51^, and are thus difficult to detect with conventional microscopy settings. To overcome these limitations in detection, longer exposure times have been employed that additionally create a bias for slow-moving mRNA and predominantly visualize mRNPs that are sequestered in large condensates such as P-bodies. It is therefore essential to use high spatial and temporal resolution imaging in living cells or image single mRNPs in fixed cells with the help of sensitive *in situ* approaches to examine the enrichment of mRNAs in biomolecular condensates.

While rapid and global translation repression following glucose withdrawal is needed to inhibit the expression of the large number of pre-stress, pro-growth genes, stress-induced transcripts can escape this repression. Earlier studies have suggested that some of the stress-induced transcripts share certain features that enable them to escape mechanisms of translation repression during stress, such as structured 5′ UTRs^29,30^ or poly(A)-rich^28^ sequences. In addition, an earlier study has reported that some stress-induced mRNAs are directed to P-bodies where they are translationally repressed, while other stress-induced mRNAs remain cytoplasmic and are efficiently translated, and that this regulation depends on the promoter sequence^25^. While specific transcripts share certain *cis-* or *trans-*regulatory sequence elements to boost their translation under global translation repression, to mount an effective stress response, cells need to activate the translation of hundreds of stress-induced transcripts that perform different biological functions and have highly diverse sequences. Since stress responses are both generic and stress-specific, a malleable translational control mechanism that goes beyond sequence-specific features could be evolutionarily advantageous.

A central unifying feature among all stress-induced transcripts is their time of production post-stress. This feature sharply distinguishes these transcripts from housekeeping transcripts that are mostly made pre-stress and transcriptionally turned off when cells encounter changes in the environment. Using an estradiol-inducible reporter that was expressed at different times relative to stress induction, we found that induction concomitant with stress led to translation whereas pre-stress induction led to translational repression. Furthermore, new mRNA, e.g. encoding ribosomal proteins, which are still produced during glucose withdrawal in wildtype cells at very low levels, or in the CARA mutant were enriched in the polysome fraction, demonstrating escape from translation repression, whereas the previously synthesized, old pool was efficiently repressed. These findings establish that the timing of mRNA production independent of mRNA sequence or promoter is a key determinant of mRNA fate in the cytoplasm. This finding is also consistent with the tight connection between promotor sequence and translation control previously observed under certain conditions ^25–27^. Thus, promotors can act as timers ensuring that mRNAs are produced at the right time to prioritize translation of newly induced mRNAs.

What drives pre-stress and post-stress mRNA to different mRNA translation fates remains an open question and a subject for further study. We can, however, postulate several hypotheses. It is likely that the prompt translational repression of pre-stress mRNA leads to major cytoplasmic reorganizations that may eventually promote the translation of the wave of newly made transcripts that begins to accumulate following stress induction. For example, translational repression of the highly abundant pre-stress mRNA may increase the number of free ribosomes/translation factors, effectively increasing the available translation capacity during stress. In addition, translational repression of pre-stress mRNA by factors such as Dhh1 may lead to sequestration of these repressors from the cytosol, thereby preventing repression of newly produced mRNA under stress. Together this may provide a competitive advantage for newly made transcripts during glucose withdrawal to get access to the translation machinery. It is also possible that newly transcribed mRNA is globally post-transcriptionally modified in a way that makes it translationally competent, e.g. with a nucleotide modification or by the binding of a protein factor. These accessory factors could preferentially interact with a component of the translation initiation machinery that is also induced in a stress-dependent manner. For example, while the 40S scanning factors eIF4A, eIF4B and Ded1 dissociate from the 5′ ends of mRNA rapidly after glucose withdrawal^19^, the helicase Dbp1 is induced under conditions of long-term nutrient deprivation and can then act as a low-performance replacement for Ded1^52^. Although this has not been shown for Dbp1 or any other translation factor following acute glucose starvation, this does not preclude the possibility that such a factor exists.

Our results suggest an elegant model of how transcription in the nucleus and translation in the cytoplasm are coordinated to ensure a timed and effective response to changes in the environment (Fig. 5). Cells turn off the expression of pre-stress mRNA predominantly encoding growth-promoting housekeeping proteins through a rapid global translation repression mechanism that keeps pre-stress mRNA in a translationally inactive state. Repression is in part mediated by decay factors to form a repressive mRNP that then also accelerates mRNA decay. Repressive mRNPs can coalesce over time into large P-body condensates but mRNA sequestration into P-bodies is not required for translational repression. At the transcriptional level, cells also mount a selective response to facilitate stress adaptation. The transcription of pre-stress, housekeeping genes is rapidly downregulated, whereas pro-stress genes are transcriptionally induced. Newly made transcripts selectively escape the translation repression, thereby ensuring that the new transcription program can be swiftly executed by the cell.

**Figure 5:**
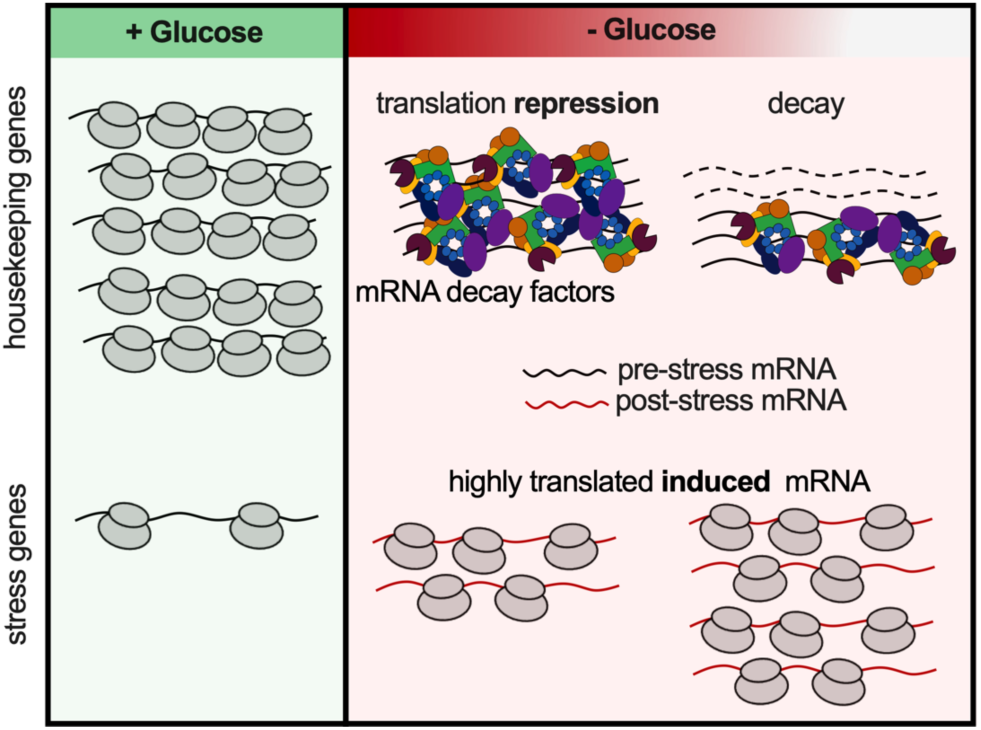
Model. Under glucose-rich conditions, a substantial fraction of yeast transcription is devoted to producing housekeeping proteins, such as ribosomal proteins and proteins involved in ribosome biogenesis, that are also highly translated under these conditions. Transcripts encoding stress-related proteins are either not produced or produced at low levels. Following stress, pre-existing housekeeping transcripts become rapidly translationally repressed by a mechanism that depends on mRNA decay factors such as Dhh1 and Pat1. While repression requires the unstructured tails of Dhh1 and thus presumably Dhh1 multimerization to form a repressive mRNP, it does not require the recruitment of these repressed mRNPs into large ribonucleoprotein condensates i.e. P-bodies. Newly made transcripts that are produced after glucose withdrawal can escape this translation repression in a time-controlled but sequence-independent mechanism and can selectively access the ribosome.

Our studies focused on the stress response that occurs in budding yeast upon acute glucose withdrawal. However, yeast cells mount similar, rapid responses to adapt to many diverse stresses including hypoxia, osmotic shock or temperature shifts. Intriguingly, an identical timing-controlled regulation is demonstrated in an accompanying paper during heat stress in budding yeast (accompanying paper by Drummond *et al*.). Furthermore, all eukaryotic cells must be able to adapt to changes in their environment, and the integrated stress response is an evolutionarily highly conserved pathway, playing critical roles for cells, tissues and organisms in development and in disease^53^. A central feature of the integrated stress response is the global inhibition of translation, predominantly affecting previously expressed housekeeping genes, accompanied by the selective and preferential translation of newly induced transcripts. It is thus tempting to speculate that a similar regulation exists in all eukaryotes and that newly synthesized transcripts can escape global translational repression under conditions where a rapid and large-scale reprogramming of gene expression is critical.

## Methods

### RNA half-life estimates (SLAM-seq)

Yeast cells (KWY165) were grown in synthetic media containing 2% glucose (SCD) and containing low uracil (10 mg/L) for two doubling times at 30 °C. To pulse label mRNA, 4-thiouracil (stock concentration 1M in DMSO) was added to the growth medium at a final concentration of 1 mM, for 120 min. To chase, cells were filtered, washed and resuspended in synthetic medium lacking 4-thiouracil and containing excess uracil (20 mM) and no 4-thiouracil, in the presence or absence of glucose. Samples were collected by filtration immediately after starting the chase (time point zero), and after 5, 10, 15, 20, 30 and 45 minutes of chase. RNA extraction was performed using hot phenol-chloroform^13^. 1 ng of spike-in RNA encoding *E. coli* transcripts were added per 10 μg of total RNA per each time point. These spike-ins were PCR amplified from extracted *E. coli* genomic DNA using primers that contain T3 promoter sequence and *in vitro* transcribed using Megascript T3 *in vitro* transcription kit. RNA alkylation was performed in 30% DMSO, as described^35^. Library preparation was performed using Quantseq for 3’ end sequencing (Lexogen, Austria). The experiment was performed in two independent replicates. Reads were mapped as described^34^ and the quantification of labeled transcripts was performed using the SLAM dunk package^54^ and normalized to spike-in levels^55^. Half-lives of transcripts were estimated by fitting the exponential decay equation as described^55^.

### mRNA Sequencing and ribosome profiling

mRNA sequencing and ribosome profiling were performed as described^56^, with some modifications. Wild type yeast cells (KWY165) or P-bodies mutants (KW7628, KWY293 and KWY8044) were incubated overnight in YPD media (containing 2% glucose) at 30°C and 150 rpm. The next day, the pre-culture was diluted in 400 ml YPD media to a starting OD_600_ of ∼0.15. The cultures are incubated until they reached an approximate OD600 of 0.6. Next, cells were filtered by a Whatman™ 0.8 µm filter, washed with 800 ml YPD media and resuspended in 400 ml of YPD media. For the starved conditions, YP media lacking glucose was used instead for the washing and resuspension step. The cultures were incubated for 5, 15 or 30 minutes after the filtration at 30°C and 150 rpm. The cells were harvested on a 0.8 µm Whatman™ cellulose filter using a vacuum pump, collected and immediately flash frozen in liquid nitrogen. Frozen cell pellets were stored at -20°C until further use. For cell lysis: Cell pellets were lysed by cryomilling with 1 ml of frozen lysis buffer (20 mM Tris-HCl pH 8.0, 140 mM KCl, 6 mM MgCl_2_, 100 μg/ml cycloheximide, 1 mM PMSF, 0.1% Tween) for 2 minutes and 30s^-1^ each. Lysates were stored at -80°C until further use. The cell lysates were thawed at 30°C for 2 minutes, cleared by centrifugation at 20,000g for 5 min, and the supernatant was transferred to a new tube. 400 µl of the sample was used for ribosome profiling and the rest of the sample was stored on ice for RNA sequencing.

For ribosome profiling, the total lysates were digested using 1.35 µl of RNAse I (10 U/μL) per 1 mg of RNA at 25°C for 1 hour. This was followed by a sucrose cushion, where 300 µl of the digested sample was carefully transferred on top of 900 µl of sucrose cushion (1M sucrose, 20 mM Tris-HCl pH 8.0, 6 mM MgCl_2_, 140 mM KCl, 100 μg/ml cycloheximide, 1 mM DTT, 2x cOmplete™ EDTA-free protease inhibitor) followed by ultracentrifugation at 70’000 rpm at 4°C for 4 hours (TLA 100.3 rotor). The ribosomal pellet was resuspended in 500 µl of 20 mM Tris-7, followed by addition of 40 µl of 20% SDS. Next, the RNA was extracted by hot phenol-chloroform extraction at 65°C and precipitated overnight using isopropanol (1:1 v/v), and 3M NaOAc pH5.2 (1:10 v/v). On the next day, the RNA pellets were washed with ice-cold 80% EtOH, and resuspended in 10 mM Tris pH 7.0. Library preparation was performed as described^56^.

For RNA sequencing: 40 µl of 20% SDS was added to the rest of the cell lysates, and the RNA was extracted by hot phenol-chloroform extraction at 65°C and precipitated overnight using isopropanol (1:1 v/v), and 3M NaOAc pH5.2 (1:10 v/v). The cell pellets were washed with ice-cold 80% EtOH. The mRNA was isolated by Poly-A selection using Invitrogen™ Oligo(dT)25 Dynabeads™. 150 μl of beads was washed twice with 100 μl binding buffer (10 mM Tris-HCl pH 7.5, 0.5M LiCl, 3.35mM EDTA). 150 μg of RNA in binding buffer was first denatured by incubation at 80°C for 2 minutes and immediately put on ice. The denatured RNA was added to the beads and incubated for 5 min at room temperature. The beads were washed twice with 100 μl washing buffer (10 mM Tris-HCl pH 7.5, 0.15 M LiCl, 1 mM EDTA), followed by resuspension in 10 mM Tris-HCl pH 7. The RNA was eluted by incubation at 80°C for 2 min, and immediately collected. The poly-A mRNA was fragmented by alkaline fragmentation in 50mM NaCO_3_ pH 9.2 and 1 mM EDTA at 95°C for 30 minutes. Samples were immediately transferred to ice and precipitated overnight using isopropanol, NaOAc pH 5.2, and 2 µl of Invitrogen™ GlycoBlue™. On the next day, the RNA pellets were washed with ice-cold 80% EtOH, and resuspended in 10 mM Tris pH 7.0 Library preparation was performed as described^56^.

### **s**miFISH

Cell culture and starvation stress induction: GFP-DCP2 (KWY2379) cells were inoculated overnight in 10 ml of SCD media containing 2% glucose. On the next day they were diluted to an OD600 of 0.2 in fresh SCD media and grown to the exponential growth phase (approx. OD600 of 0.7). Next, cells were harvested on a Whatman™ cellulose filter using a vacuum filtration pump, resuspended in SC media and incubated for the specified time of starvation stress. Unstressed cells were treated equally except that media switch happened into SCD media containing 2% glucose.

Cell fixation and spheroplasting: The experimental procedures were carried out according to an optimized smFISH protocol^57^ with slight adaptations. Briefly, cells were fixed with 4% PFA, washed twice with ice-cold buffer B (1.2 M Sorbitol, 100 mM KHPO4 pH 7.5, 4°C) and transferred to a 5x concanavalin A (stock: 1 mg/ml) coated and air-dried 96-well plate. After a short spin of the plate, the cells walls were digested by adding a spheroplasting solution (1.2 M Sorbitol, 0.1 M KHPO4 pH 7.5, 20 mM β-Mercaptoethanol) containing 1% zymolase 20T (stock: 5 mg/ml diluted in 1.2 M sorbitol) and 10% vanadyl ribonuclease complex (VRC, stock 200 mM) and incubating at 30°C for 15 minutes. The zymolase was removed by washing cells once with Buffer B containing 10% VRC.

Probes, prehybridization and hybridization: smiFISH primary and labeled FLAP probes were purchased from Microsynth. Next, a pre-hybridization buffer (10% deionized formamide, 2x saline-sodium citrate buffer SSC, 10% VRC) was added to the cells. The hybridization buffer was prepared in advance by heating up a 1:1 mixture of a mRNA binding oligo (25µM Stock) and its labeled FLAP (25 µM Stock) to 95°C for 3 minutes for annealing. In case of the negative control (+/-) only the secondary labeled FLAP was mixed 1:1 with TE buffer. The remaining components of the hybridization buffer (0.19 mg/ml yeast transfer RNA, 0.19 mg/ml salmon-sperm DNA, 10% deionized formamide, 5 mM NaHPO4 pH 7, 2x saline-sodium citrate buffer SSC, 2 mg/ml acetylated BSA, 10% VRC) were added to the Oligo-FLAP mix once it had cooled down to room temperature. 1 µl of probe mix and 104 µl of the hybridization buffer were mixed for each sample. Out of this mixture, 80 µl were added to each sample. Depending on the mRNA which was targeted, the probe mixture was diluted in a range from 1x to 1:10x in TE buffer (pH 8.0, 10 mM Tris-HCl containing EDTA·Na2). The dilution leading to the best signal-to-noise ratio was determined by titration. The 96-well plate was sealed with a Greiner Bio-one BREATHseal™ Sealer to reduce evaporation and an aluminum foil on top to protect the samples from light. Cells were incubated overnight at 37°C. On the next day, cells were transferred to a freshly coated 96-well plate and washed once with 1x PBS/DAPI (1:4000) and once with 1x PBS.

Imaging: Imaging was performed at room temperature on an inverted epifluorescence microscope (Nikon Ti) equipped with a Spectra X LED light source and a Hamamatsu Flash 4.0 sCMOS camera using a PlanApo 100× NA 1.4 oil-immersion objective and the NIS Elements software (Version 5.21.03). Of each sample, BF, RFP (ExW=555nm), GFP (ExW=475nm) and DAPI (ExW=395nm) images were acquired with the same settings in 11 z-planes were on a total of 4 µm distance, with the following exposure times: RFP=400ms, GFP=300ms, DAPI=300ms.

Image analysis: The analysis was performed using in-house scripts and consisted of three main steps: 1. Producing maximum projections 3 center slices of the acquired image z-stacks and splitting the channels (mRNAs, P-bodies, BF) into different files. 2. Detection of the single mRNA localizations using a ThunderSTORM^58^(Fiji^59^). For this localization step, the intensity threshold was chosen based on the negative control images, where no mRNAs should be detected. ThunderSTORM outputs a file containing the XY coordinates of all the detected mRNAs. This matrix was used as an input for a custom script written in Matlab (MathWorks Inc. Version R2022b) that performed the rest of the analysis. 3. The P-bodies were segmented based on an intensity threshold and polygons enclosing each P-body were obtained. The same threshold (600 a.u) was used for the PB detection, independent of the mRNA candidate, because this signal was not expected to vary significantly among samples which were starved for equal durations. This threshold was defined for GFP-DCP2 tagged PBs in 30-minute starved *S. cerevisiae* cells. Then, the intensity of single mRNAs outside P-bodies was calculated. The brightness of individual localizations (corresponding to individual mRNAs molecules) and their cumulative intensities were obtained and subtracted from their corresponding local background. For that, the intensity of all pixels (I_1_) was summed within a 5x5 pixels area (A_1_) around the localization position (x, y) of the fluorophore. The same operation was then performed for a larger, 7x7 pixels area (A_2_) around the same location, to obtain I_2_. The local background was calculated from the pixel values between the smaller and larger regions of interest and subtracted from I_1_ accordingly^60^:

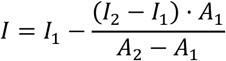

The single mRNAs intensity distribution for each individual cell was fitted with a Gaussian function to obtain the average single mRNA intensity^60^. Then, the same procedure was applied to obtain the intensity of the mRNA channel in the segmented P-body regions. In this case, A_1_ will correspond to the segmented ROI of the P-body, and A_2_ will be an expanded polygon encompassing local background pixels. Note that the polygons until the proportion of the number of A_2_/A_1_ pixels is similar to the one used for calculating the intensity of single mRNAS (7x7/5x5). We then divided the total normalized intensity inside the P-body regions by the average single mRNA intensity, to estimate the number of mRNAs inside the P-bodies.

### Reporter induction and polysome profiling

Yeast cells (KWY7227) were grown in synthetic media containing 2% glucose (SCD) for at least two doubling times to log phase at 30 °C. For induction without stress, beta-estradiol (stock concentration 400 uM in DMSO) was added to a final concentration of 400 nM, and incubated for 30 minutes followed by harvesting using filtration and snap-freezing in liquid nitrogen. For pre-stress induction, beta-estradiol was added to log-phase yeast cells growing in SCD to a final concentration of 400 nM, and incubated for 30 minutes at at 30 °C. Cells were harvested by filtration, washed and resuspended in synthetic medium lacking glucose (SC) and lacking beta-estradiol for 30 minutes followed by harvesting using filtration and snap-freezing in liquid nitrogen. For post-stress induction, log-phase yeast cells growing in SCD were harvested by filtration, washed in synthetic medium lacking glucose and resuspended in synthetic medium lacking glucose and containing beta-estradiol at final concentration of 400 nM, for 30 minutes followed by harvesting using filtration and snap-freezing in liquid nitrogen. Cell pellets were lysed by cryomilling with 1 ml of frozen lysis buffer (20 mM Tris-HCl pH 8.0, 140 mM KCl, 6 mM MgCl2, 100 μg/ml cycloheximide, 1 mM PMSF, 0.1% Tween) for 2 minutes and 30s^-1^ each. The cell lysates were thawed at 30°C for 2 minutes, and cleared by centrifugation at 20,000g for 5 min, and the supernatant was transferred to a new tube. The total lysate was loaded into 10-50% sucrose gradient followed by centrifugation for 2.5 hr at 35,000 rpm and 4°C, followed by gradient fractionation and absorbance measurement at 260 nm, and the polysomes fraction was pooled together. RNA was extracted from the total lysate or from the polysome fraction, as described^56^. Library preparation was performed using Nextflex rapid directional RNA sequencing kits.

### mRNA 4-TU labeling and polysome profiling

Yeast cells (KWY11376) were grown in synthetic dropout medium lacking histidine and uracil, and containing 2% glucose (SCD -HIS -URA) for at least two doubling times to log phase at 30 °C. For labeling under glucose-rich conditions, 4-thiouracil (final concentration of 1 mM), or beta-estradiol (final concentration of 400 nM) was added followed by incubation for 30 minutes. Cells were harvested by filtration and snap-frozen in liquid nitrogen, or transferred into (SC -HIS -URA) medium lacking glucose, and lacking 4-thiouracil or beta-estradiol, for 30 minutes before harvesting (pre-stress induction). For glucose withdrawal and mRNA labeling, yeast cells growing in glucose-rich conditions (SCD -HIS -URA) were harvested by filtration, washed and resuspended in (SC -HIS -URA) lacking glucose and containing 4-thiouracil at a final concentration of 1 mM or beta-estradiol at a final concentration of 400 nM, for 30 minutes. Cells were harvested by filtration and snap-frozen in liquid nitrogen. Cell were lysed by cryomilling, followed by polysome fractionation and extraction of RNA from the total lysate or from the polysome fraction, as described^56^. Cell pellets were lysed by cryomilling with 1 ml of frozen lysis buffer (20 mM Tris-HCl pH 8.0, 140 mM KCl, 6 mM MgCl2, 100 μg/ml cycloheximide, 1 mM PMSF, 0.1% Tween) for 2 minutes and 30s^-1^ each. The cell lysates were thawed at 30°C for 2 minutes, and cleared by centrifugation at 20,000g for 5 min, and the supernatant was transferred to a new tube. RNA was loaded into 10-50% sucrose gradient followed by centrifugation for 2.5 hr at 35,000 rpm and 4°C, followed by gradient fractionation and absorbance measurement at 260 nm, and the polysome fraction was pooled together. RNA was extracted from the total lysate or from the polysome fraction, as described^56^. RNA alkylation was performed in 30% DMSO, as described^35^. Library preparation was performed using Nextflex rapid directional RNA sequencing kits. Library preparation was performed using Quantseq for 3’ end sequencing (Lexogen, Austria). Similar experiments were performed using KWY12663 and KWY12664 grown in complete synthetic media for mRNA labeling using 4-TU during glucose starvation.

## Acknowledgements

We thank members of the Weis lab for discussions and critical comments on the manuscript, and Allan Drummond for sharing of results prior to publication. We are grateful to Christophe Carles and Herbert Tschochner for their help and the gift of the CARA strains. This work was supported by an EMBO long-term fellowship to S.H. (ALTF 290-2014), and an ETH postdoctoral fellowship to P.G.G. (22-1 FEL-06). This work was also supported by grants from the Swiss National Science Foundation to K.W. (TMAG-3_209354 and CRSII5_193740).

## Author Contributions

Conceptualization, M.Z. and K.W.; Methodology, M.Z., S.H., P.G.G. and K.W.; Investigation, M.Z., A.S. and P.G.G.; Formal analysis, M.Z., A.S. and P.G.G.; Writing – Original Draft, M.Z, S.K. and K.W.; Writing – Review & Editing, M.Z., A.S., P.G.G., S.K. and K.W.; Funding Acquisition, P.G.G and K.W.; Resources, K.W.; Supervision, M.Z. and K.W.

## List of strains

|  |  |
| --- | --- |
| KWY165 | W303 MAT A his3-11,15 ura3-1 leu2-3 trp1-1 |
| KWY293 | W303 MAT A dhh1Δ::KAN |
| KWY7628 | W303 MAT A his3-11, 15ura3-1 leu2-3 trp-1 Dhh1:48-425 yEGFP::HIS |
| KWY8044 | W303 MAT A pat1Δ::Kan |
| KWY2379 | W303 MAT A DCP2::GFP HIS5 |
| KWY7227 | BY4741 MAT A his3_1 leu2_0 met15_0 ura3_0 Nup60-3xmKate2::CaURA3 pNH604-pGPD1-LexA-B112::HIS Chromosome2(CHS3-SCO2 intergenic region)::p(LexA(4)-pCYC1min)_3xGST-V5_24xPP7sl_Tcyc1_NatNT2 pMET25-PP7CP-yEGFP::LEU2 Dcp2-mCherry::KAN |
| KWY11376 | W303 MAT A his3-11,15::HIS3 ura3-1::URA3 leu2-3 trp1-1 pACT1(-1-520)-LexA-ER-haB112-tCYC1 insul-(lexAbox)x4-pMinCYC1-Citrine-tCYC1 |
| KWY12663 | MATα ade2-101 his3-200 leu2-1 lys2-801 trp1-63 ura3-52 rrn3::his5 rpa43::kan pGEN-RRN3-A43(TRP1) |
| KWY12664 | MATα ade2-101 his3-200 leu2-1 lys2-801 trp1-63 ura3-52 rrn3::his5 |

---

**Supplementary Figure 1:**
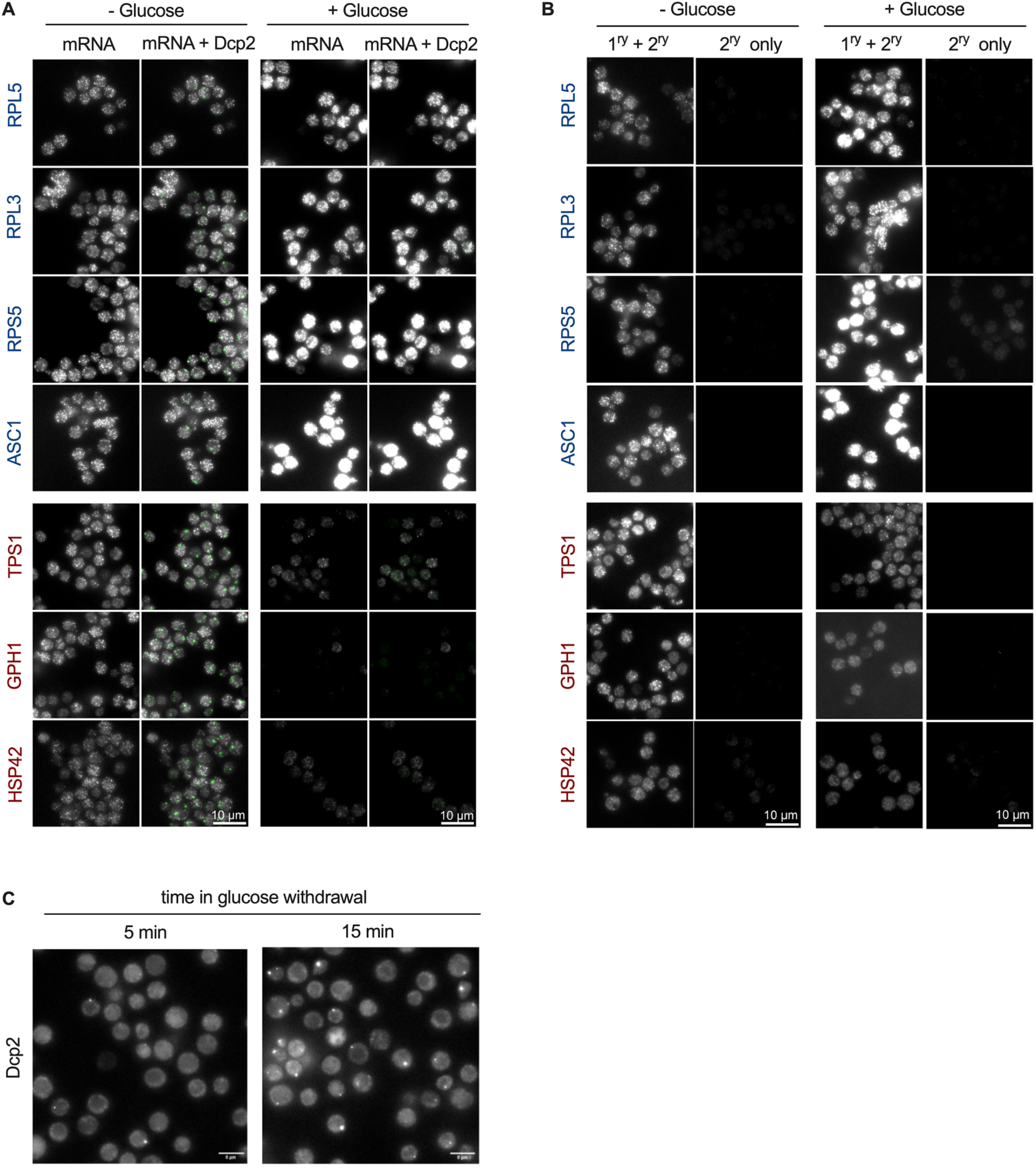
smiFISH analysis of target mRNAs under glucose-rich or glucose withdrawal conditions. **A.** smiFISH analysis under glucose-rich or glucose withdrawal conditions. Yeast cells expressing GFP tagged Dcp2 at the endogenous locus (as marker of P-bodies) were grown to log phase in glucose-rich medium. Glucose withdrawal was performed by shifting yeast cells to glucose-free medium by rapid filtration and incubation for 30 minutes. 27-48 primary probes complementary to the target mRNA sequence were used per candidate mRNA. The primary probes containing the flap sequence were pre-hybridized with an oligo double labeled with a TAMRA dye (secondary probe) complementary to the flap-sequence of the primary probes, before being hybridized *in situ* to the target mRNA. Translationally downregulated mRNAs following glucose withdrawal are colored in blue, and translationally upregulated mRNA following glucose withdrawal are colored in red. **B.** Secondary probes were used in the absence of primary probes using identical hybridization and imaging conditions as in A. **C.** Analysis of P-body formation after 5 or 15 minutes of glucose withdrawal. Yeast cells expressing GFP tagged Dcp2 at the endogenous locus (as marker of P-bodies) were grown to log phase in glucose-rich medium. Glucose withdrawal was performed by shifting yeast cells to glucose-free medium by rapid filtration and incubation for 5 or 15 minutes.

**Supplementary Figure 2:**
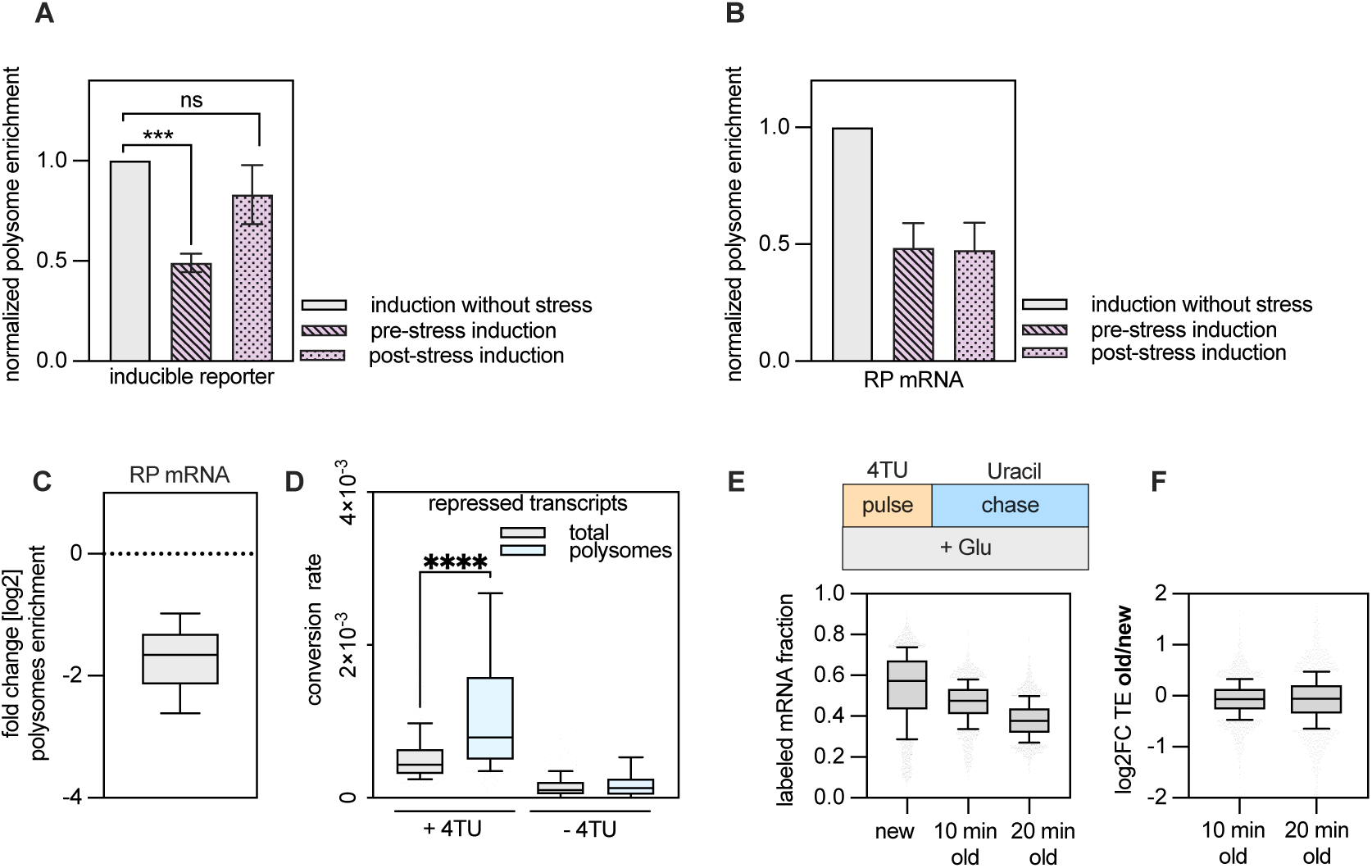
The timing of mRNA production determines translation efficiency. **A.** Analysis of polysome enrichment of an estradiol-inducible reporter mRNA encoding citrine following glucose withdrawal after pre- or post-stress mRNA induction, relative to polysome enrichment of the reporter mRNA induction under glucose-rich conditions. The enrichment of mRNA in polysomes was quantified by normalizing mRNA levels in the polysome fraction to its total levels in the lysate. Bars represent the median of two replicates, and the error bars represent the 95% confidence interval. **B.** Analysis of polysome enrichment of mRNA encoding ribosomal proteins following glucose withdrawal after pre- or post-stress mRNA induction of the reporter mRNA encoding citrine protein, relative to polysome enrichment of mRNA encoding ribosomal proteins under glucose-rich conditions where the reporter mRNA encoding citrine protein was also induced. The enrichment of mRNA in the polysomes was quantified by normalizing mRNA levels in the polysome fraction to its total levels in the lysate. Bars represent the median of two replicates, and the error bars represent the 95% confidence interval. **C.** Box plot showing the change in polysome enrichment (in log_2_) of old and newly made RP mRNA following glucose withdrawal and mRNA labeling using 4-TU for 30 minutes, compared to 4-TU labeling of transcripts under glucose-rich conditions for 30 minutes. Polysome enrichment was calculated by normalizing mRNA levels in the polysome fractions versus total mRNA levels. All reads (labeled and unlabeled) were included in the analysis. The experiment was performed in two replicates. **D.** Analysis of polysome enrichment of newly made transcripts that show at least 2-fold decrease in translation efficiency following 30 minutes of glucose withdrawal. To label newly made transcripts after glucose withdrawal, yeast cells were shifted from glucose-rich medium to glucose-free medium lacking uracil but containing 4-TU for 30 minutes before harvesting. To analyze the level of mRNA translation, cell lysates were fractionated using a sucrose gradient followed by polysome isolation. Labeled transcripts in the total lysate or in the polysomes were quantified based on the calculated T>C conversion rate. The experiment was done in two replicates. To control for background T>C conversion, the experiment was repeated without addition of 4-TU. **E.** Analysis of the fraction of labeled mRNA following 10 minutes of 4-TU pulse labeling under glucose rich conditions (“New” mRNA), as well as 10 or 20 minutes after chasing with excess uracil under glucose rich conditions (“old” mRNA). Labeled transcripts were quantified based on the detection of T>C conversions. **F.** Comparison of translation efficiency change of old (10 or 20 minutes after chase) versus new (after 10 minutes of pulse labeling with 4-TU) under glucose rich conditions. Translation efficiency was calculated by normalizing the levels of labeled transcripts in the polysomes fraction to the total lysate levels.

